# Integrated tissue proteomics and lipidomics from ovine organs suggests novel functional implications for fatty acids and lipid mediators

**DOI:** 10.1101/2025.10.28.685055

**Authors:** Gerhard Hagn, Florien Jenner, Andrea Bileck, Sinan Gültekin, Mats Hamberg, Thomas Mohr, Christopher Gerner, Iris Gerner

**Author notes:** Correspondence: Iris Gerner,; Christopher Gerner. Contributed equally (Gerhard Hagn, Florien Jenner, Christopher Gerner, Iris Gerner).

## Abstract

Lipid mediators are potent biomolecules that may help to gain specific information about biological processes. However, due to incomplete functional annotations and hence interpretive challenges these molecules are hardly considered in molecular profiling experiments. We hypothesized that correlating lipid mediators extracted from tissues with corresponding proteomics data and the corresponding Gene Ontology terms characteristic of specific tissue types could enable functional annotation of these lipids. Thirteen organs and tissues from sheep (*Ovis aries*) were analyzed using mass spectrometry-based untargeted proteomics and lipidomics. In total, 4717 proteins and 166 free fatty acids, oxylipins, lysolipids, endocannabinoids, and bile acids were catalogued. This included the previously uncharacterized oxylipin 4-hydroxy-eicosatetraenoic acid (4-HETE), whose identity was confirmed by chemical synthesis. Co-expression and clustering analyses validated our hypothesis, successfully reproducing known and suggesting novel lipid mediator functions. Tissue levels of polyunsaturated fatty acids (PUFAs) correlated with pro-resolving mediators and, unexpectedly, with the cellular protein synthesis and folding machinery. These findings suggest that PUFAs and their derivatives support protein synthesis fidelity and efferocytosis, both essential contributors for the effective resolution of inflammation.

Graphical abstract:
The Figure was partly generated using Servier Medical Art, provided by Servier, licensed under a Creative Commons Attribution 3.0 Unported license.

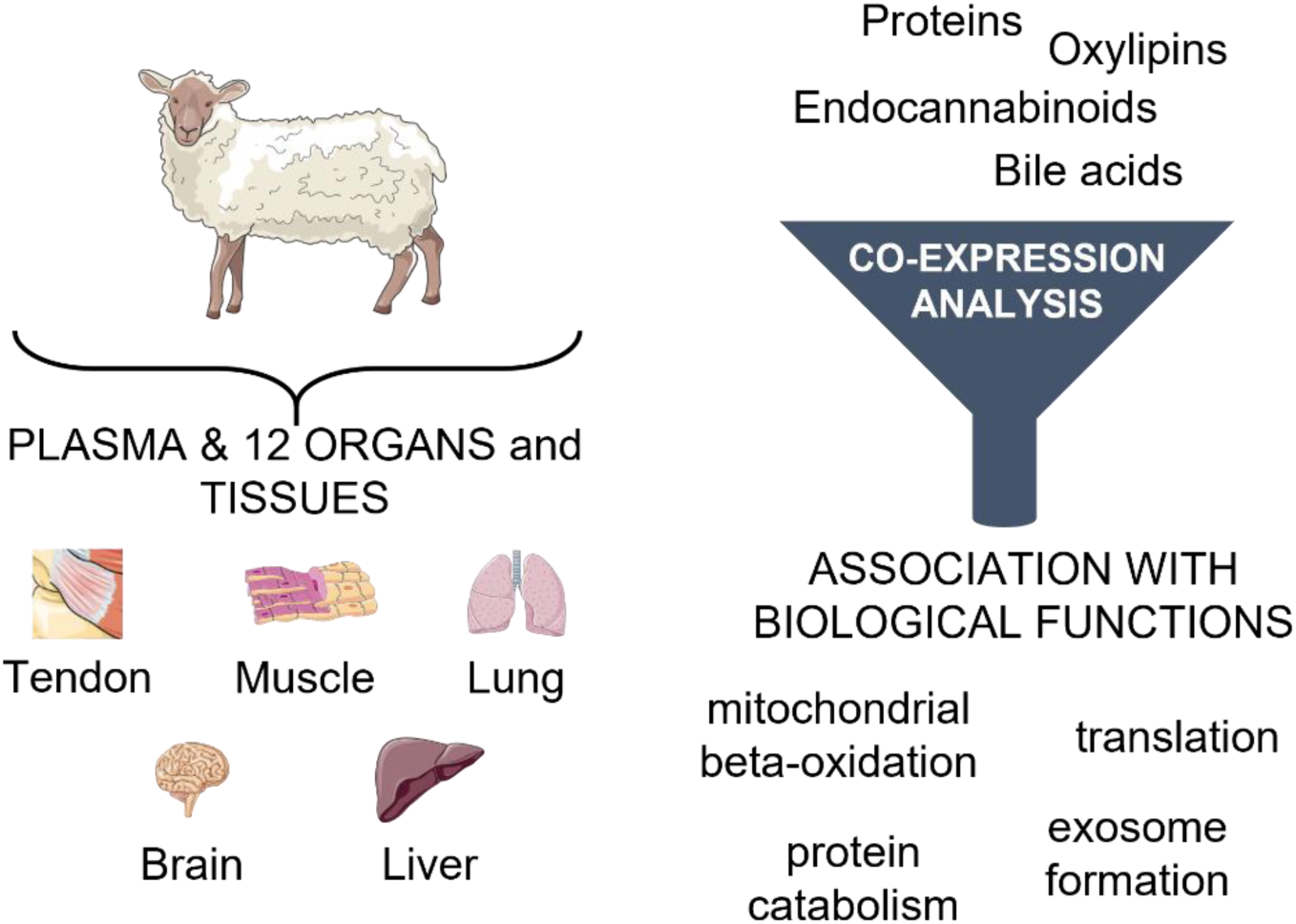

## Introduction

Metabolic activities are known to be intricately linked with signaling and transcriptional networks that regulate cellular activities [1–3]. The metabolic profiles of cells change in response to substrate availability and throughout their lifespan to support their functional roles as they differentiate, age, or encounter stressors [4]. For instance, the metabolic activities in proliferating cells differ fundamentally from those in non-proliferating cells [2; 5]. The tightly coordinated interplay between cellular physiology and metabolism is emerging as a fundamental determinant of tissue health and longevity and deserves further investigation [6–9].

Lipid mediators are metabolites which may also act as signaling molecules, binding as ligands to specific receptors [5; 10-12]. Already more than three decades ago the term “oxylipins” was defined upon recognizing that a complex enzymatic system may exist which forms specific lipid mediators from polyunsaturated fatty acids (PUFAs) [13]. This connection suggests that lipid mediators may serve as indicators for the actual exertion of specific processes. We have already observed characteristic alterations of lipid mediators in plasma samples in the course of an acute inflammatory challenge [14]. This motivated us to investigate the relation of lipid mediators with physiologic processes in a systematic fashion. The present study was designed accordingly and is based on two assumptions. First, different types of organs and tissues are associated with the exertion of specific and well-known biological processes. Indeed, different tissues also exhibit distinct metabolic profiles [15]. In adult mammals, physiological cell functions, but also the rate of cell proliferation and turnover, vary significantly between tissues, depending on each tissue’s specific function and environmental cues [16]. The most characteristic biological processes associated with a given organ or tissue are strongly associated with the proteome, and accessible from experimental data *via* the gene ontology classification system [17]. The second assumption of this study is that the abundance of a given lipid mediator may correlate with the exertion of a corresponding biological process. As a consequence, a correlation analysis of biological processes retrieved from proteomics data with lipid mediators retrieved from lipidomics analyses across different organs and tissues should allow us to relate the abundance of lipids with the probability for the exertion of biological processes.

Actually, data on relevant metabolites and their functional implications remain scarce [18–20] and metabolomic studies have predominantly focused on tissues involved in energetically demanding tasks, such as physical work (e.g., skeletal and cardiac muscles), neurological processes (brain), or vital physiological functions (liver, kidney), with little attention on bradytrophic tissues composed of largely quiescent cells such as cartilage and tendon. Therefore, this study aims to determine the lipid mediator fingerprints of plasma and 12 organs and tissues with differing metabolic activities (brain, heart, kidney cortex, renal medulla, liver, lung, muscle, nerve, skin, spleen, tendon, cartilage). The association of the occurrence of lipid mediators with biological functions was investigated using an integrated mass spectrometry-based lipidomics and proteomics approach. We opted to utilize sheep (*Ovis aries*), a well-established large animal model, for this study. In contrast to small animal models such as mice, they are a long-lived species, and their body size, weight, and organ dimensions closely resemble those of humans [21; 22]. Already as early as in 1982, fundamental studies on the regulatory effects of oxylipins on the vascular tone have been performed in sheep [23]. However, while previous efforts have focused on the global analysis of gene expression across multiple ovine tissues, mass spectrometry (MS)-based proteome and metabolome data for the characterization of (ovine) tissues and organs with respect to molecules relevant for immune modulation hardly exist [24]. We have established robust and reliable oxylipin and proteome profiling methods and have successfully combined these techniques for biomedical studies [25–27]. The poor functional annotation of sheep proteins by gene ontology was compensated using gene ontology annotations of human orthologues, previous work demonstrates this to work well [21; 28]. This facilitated the analysis of co-expression networks and clustering of the combined dataset as described previously [29]. The applicability of this unique bioinformatical approach was presently tested for the first time for proteins and oxylipins. This feasibility study resulted in numerous potential functional annotations of lipids which may help to better understand the physiological roles and potential disease implications of these important molecules.

## Materials and methods

### Sample acquisition and Ethics

Organs and tissue were collected from three (n=3), healthy, skeletally mature (2-4 years of age), female sheep (Merino) with a body weight between 70 and 90 kg which had been euthanized for reasons unrelated to this study. Based on the “Good Scientific Practice, Ethics in Science and Research regulation” implemented at the University of Veterinary Medicine Vienna, the Institutional Ethics Committee and the Austrian Federal Ministry of Educations Science and Research, *in vitro* studies do not require approval if they are performed using tissues/organs which were obtained either solely for diagnostic or therapeutic purposes or in the course of other institutionally and nationally approved experiments.

Blood (5 mL) was collected from the jugular vein into ethylenediaminetetraacetic acid (EDTA) tubes and plasma was obtained *via* centrifugation (10 min, 4 °C, 2000 g) and stored at −80 °C until further analysis. In addition, samples were taken from twelve organs and tissues, the brain (caudal aspect of cerebrum), heart (apex), kidney cortex and renal medulla, liver (caudal lobe), lung (accessory lobe), muscle (m. gastrocnemius), nerve (brachial plexus), skin (lateral knee region), spleen, tendon (superficial digital flexor tendon of one front leg, mid tendon region) and cartilage (from the medial femoral condyle). For proteomics, about 5 – 10 mg of each organ/tissue were collected, whereas about 50 – 100 mg were collected for the lipid analysis. To remove blood cells as completely as possible, the obtained organ/tissue pieces were rigorously rinsed with ice cold phosphate-buffered saline (PBS) until no residual blood was visible on the sample surfaces. Care was taken that the organs/tissues were exsanguinated to the greatest possible extent, while the blood vessels and capillaries were not flushed. After washing, the tissue samples for proteomics analysis were immediately snap frozen in liquid nitrogen and stored at −80 °C. The samples for the oxylipin/fatty acid analysis were transferred into 15 mL Falcon^TM^ tubes containing 1 mL of ice-cold ethanol (EtOH) and stored at −20 °C.

A detailed description of the following paragraphs on proteomics and lipidomics sample preparation, data acquisition and processing can be found in the *Supporting Information SI1*.

### Proteomics Sample Preparation

For untargeted tissue and plasma proteomics analysis, frozen tissue samples were transferred into 15 mL Falcon^TM^ tubes containing 150 µL of lysis buffer (8 M urea, 50 mM triethylammonium bicarbonate (TEAB), 5 % sodium dodecyl sulfate (SDS)) and homogenized using an ultrasound sonication homogenizer. After centrifugation at 5000 g and room temperature (RT) for 5 min, the supernatant was transferred into fresh tubes. Ovine plasma was diluted 1:20 using lysis buffer, and tissue and plasma samples were heated for 5 min at 95 °C and 300 rpm under constant shaking. After cooling to RT, the protein concentration was determined *via* a bicinchoninic acid (BCA) assay. Aliquots of plasma and tissue samples containing 20 µg protein were used for enzymatic protein digestion according to the ProtiFi S-Trap^TM^ protocol as described [30].

### Proteomics Data Acquisition

Mass spectrometric analysis was performed with a timsTOF Pro (Bruker, Germany), in the Parallel Accumulation-Serial Fragmentation (PASEF) mode. The scan range was 100 to 1700 m/z for MS and MS/MS spectra recording and the 1/k0 scan range was set to 0.60 – 1.60 V*s cm^-2^ resulting in a ramp time of 100 ms to achieve trapped ion mobility separation.

### Proteomics Data Processing

Protein identification was performed *via* MaxQuant [31] (version 1.6.17.0) employing the Andromeda search engine against the UniProt Database [32] (version 11/2021, 20’375 entries). Search parameters were set as previously described [33]. Proteins with at least 60 % identification rate in at least one group were considered for analysis. A two-sided Student’s t-test with S0 = 0.1 and an FDR cut-off of 0.05 was applied for identifying multiple testing-corrected significantly regulated proteins.

### Lipidomics Sample Preparation

For untargeted tissue and plasma analysis of oxylipins and fatty acids, 400 mg EDTA-anticoagulated plasma was thawed on ice and added to a 15 mL Falcon^TM^ tube containing ice-cold EtOH (1.6 mL, abs. 99.9 %, AustrAlco) and an internal standard mixture (Supplementary Table S1). All standards, including those indicated in Supplementary Table S2, were purchased by Cayman Chemical, Michigan, USA. The Falcon^TM^ tubes containing tissue samples and ethanol were transferred from −20°C to ice, homogenized using an ultrasound sonication homogenizer and stored again at −20°C overnight. On the following day, LC-MS grade water and the same internal standard mix that was used for the plasma samples were added, and all plasma and tissue samples were centrifuged. The supernatant was transferred, and the ethanol was evaporated. Solid phase extraction of tissue and plasma samples was performed as previously described [34].

### Lipidomics Data Acquisition

For the LC-MS/MS analyses, a Vanquish™ UHPLC system (Thermo Fisher Scientific™, Vienna, Austria) was coupled to a high-resolution quadrupole orbitrap mass spectrometer (Thermo Fisher Scientific™ Q Exactive™ HF hybrid quadrupole orbitrap mass spectrometer). Therefore, the Vanquish™ ultra-high performance LC (UHPLC) system was equipped with a reversed-phase Kinetex^®^ XB-C18 column (2.6 µm XB-C18, 100 Å, LC column 150 × 2.1 mm, Torrance, CA, USA) to achieve analyte separation as previously described [14]. Thirty-three *m/z* values specific for oxylipins and their precursor fatty acids from an inclusion list were preferentially selected for fragmentation (Supplementary Table S3).

### Lipidomics Data Processing

For data analysis, raw files generated by the Q Exactive™ HF high-resolution mass spectrometer were evaluated using the TraceFinder software (version 4.1; Thermo Fisher Scientific™, Vienna, Austria) making use of measurements of purchased chemical standards (Supplementary Table S2). Additionally, MS/MS fragmentation spectra were evaluated referring to reference spectra of the commercially available standards or to reference spectra from the LIPID MAPS repository library [35] using the Xcalibur™ Qual Browser software (version 4.1.31.9; Thermo Fisher Scientific™, Vienna, Austria). Coeluting isomers were reported giving the names of both analytes, e.g., 12-HETE/8-HETE, as suggested by the oxylipin community [36]. The TraceFinder software (version 4.1; Thermo Fisher Scientific™, Vienna, Austria) was used for relative quantification allowing a mass deviation of 5 ppm as described before [14]. The resulting peak areas of each analyte with a S/N greater than 10 were exported and read using the R software package (version 4.2.0) [37]. The peak areas were log2-transformed and analyte peak areas were normalized to the internal standards to correct for variances arising from sample extraction and LC-MS/MS analysis. Therefore, the log2-transformed mean peak area of the internal standards was subtracted from the log2-transformed analyte areas. The peak areas of the ovine plasma, organ and tissue samples were also normalized to the weight. Only molecules independently identified in all replicates of at least one tissue type were included in the statistical analysis. As several molecules turned out to occur at the limit of detection in other tissue types, data imputation was used to enable subsequent statistical analysis. To enable an imputation of missing values using the minProb function of the imputeLCMD package (version 2.1), 20 was added to the log2-transformed normalized areas to obtain values similar to proteomics label free quantification (LFQ) values.

### Data visualization

The extracted ion chromatogram was generated using the Xcalibur™ Qual Browser software (version 4.1.31.9; Thermo Fisher Scientific™, Vienna, Austria). Further data visualization was performed using the ggplot2 R package [38].

### Data Sharing Statement

Data derived from the lipidomics analysis are available at the NIH Common Fund’s National Metabolomics Data Repository (NMDR) website, the Metabolomics Workbench, https://www.metabolomicsworkbench.org (accessed on 29 May 2025) [39], where they have been assigned to the following studies: Study ID ST003163 and ST003164. The DOI for this project (PR001968) is: http://dx.doi.org/10.21228/M8BF0G.

The mass spectrometry proteomics data have been deposited to the ProteomeXchange Consortium *via* the PRIDE [40] partner repository with the dataset identifiers PXD064992 and PXD065008.

### Synthesis of the 4-HETE standard

A detailed description of the synthesis of the 4-HETE and the GC-MS analysis results are provided in *Supporting Information SI1*.

## Results

### Identification and relative quantification of lipid mediators in ovine plasma and 12 organ and tissue types

Taking account for the short half-life of many oxylipins, tissue samples were collected from anesthetized sheep and immediately snap frozen. The application of solid phase extraction to ethanol extracts of the homogenized sheep tissue samples allowed us to enrich various lipid mediators comprising fatty acids, oxylipins, lysolipids, endocannabinoids and bile acids. Using high-resolution mass spectrometry and 95 analytical standards, 166 distinct molecules were reproducibly identified in all three replicates of at least one organ or tissue type (Supplementary Table S2, the level of identification is indicated for each molecule). For comparative analysis, the determined abundance values (area under the curve values) were normalized using a set of internal standards.

Additional 42 molecules, including the previously undefined oxylipin 4-hydroxy-eicosatetraenoic acid (4-HETE, Figure 1), were identified as oxylipins based on the sum formula and fragment spectra. Due to high structural ambiguity of oxylipin isobars, no unambiguous structures were yet assigned to these molecules. These oxylipins were presently designated according to the formula “molecular mass”_”chromatographic retention time”. The applied analytical procedure is exemplified for the previously undefined oxylipin 4-hydroxy-eicosatetraenoic acid (4-HETE, Figure 1). The extracted ion chromatogram illustrates the chromatographic separation of various hydroxy-eicosatetraenoic acids (HETEs) and isobaric epoxy-eicosatrienoic acids, which differ with regard to the position of the hydroxy- or epoxy-group within the molecule. The identification of 5-HETE, 8-HETE, 9-HETE, 11-HETE, 12-HETE and 15-HETE as well as 11,12-epoxy-eicosatrienoic acid (EET) and 14,15-EET was successfully verified by commercially available analytical standards, as they displayed the same chromatographic retention times, exact molecular mass, isotopic pattern and fragmentation patterns. However, no analytical standard was available for 4-HETE (Figure 1), presently identified in heart, kidney cortex, liver, lung, muscle and spleen tissue. This oxylipin is not yet listed in LIPID MAPS, a database of well characterized lipids with largely known biochemistry [35]. In the first step, the present identification of 4-HETE was based on the exact molecular mass and the assignment of specific fragment ions as indicated in Figure 1. The MS/MS fragmentation pattern, characterized by the specific fragment of m/z = 101.024, is a well-known marker of oxylipins with a functional group at position C4 such as the docosahexaenoic acid (DHA)-derived products 4-hydroxy-docosahexaenoic acid (HDoHE or HDHA) (*Supporting Information SI1* - Supplementary Figure 1) or 4-F4t-Neuroprostane (LIPID MAPS – Standards Spectra Library; LMFA04010524). Subsequently, we were able to verify this identification *via* chemical synthesis and LC-MS analysis (retention time difference of less than 0.01 min between ovine tissue samples and synthetic standard) thereof. The 4-HETE synthesis protocol and analytical verification *via* GC-MS is described in detail in the *Supporting Information SI1*.

**Figure 1:**
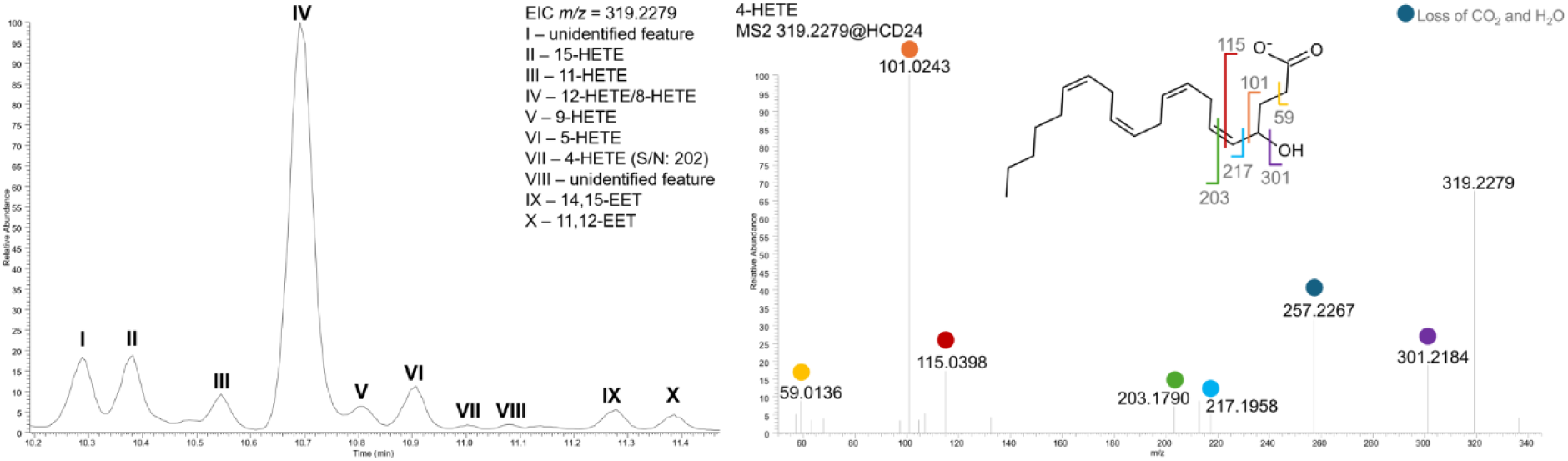
Identification of the previously uncharacterized oxylipin 4-HETE by LC-MS/MS. A: Extracted ion chromatogram (EIC) for m/z = 319.2279 obtained from a lung sample. B: MS2 spectrum obtained for peak VII (S/N: 202) indicated in the EIC and the assignment of the identifying molecular fragments according to the molecular structure of 4-HETE.

Evidently, additional effort needs to be spent for the characterization of potential physiologic effects of this previously undescribed oxylipin.

### Lipid mediators display a tissue-specific expression pattern

While some molecules such as deoxycholic acid showed low specificity with regard to organ and tissue types (Figure 2A), others were exclusively identified in one or a few thereof. This is exemplified for DHA isoform I, representing DHA with one trans-configured double bond as described previously [34], which was detected only in plasma (Figure 2B).

**Figure 2:**
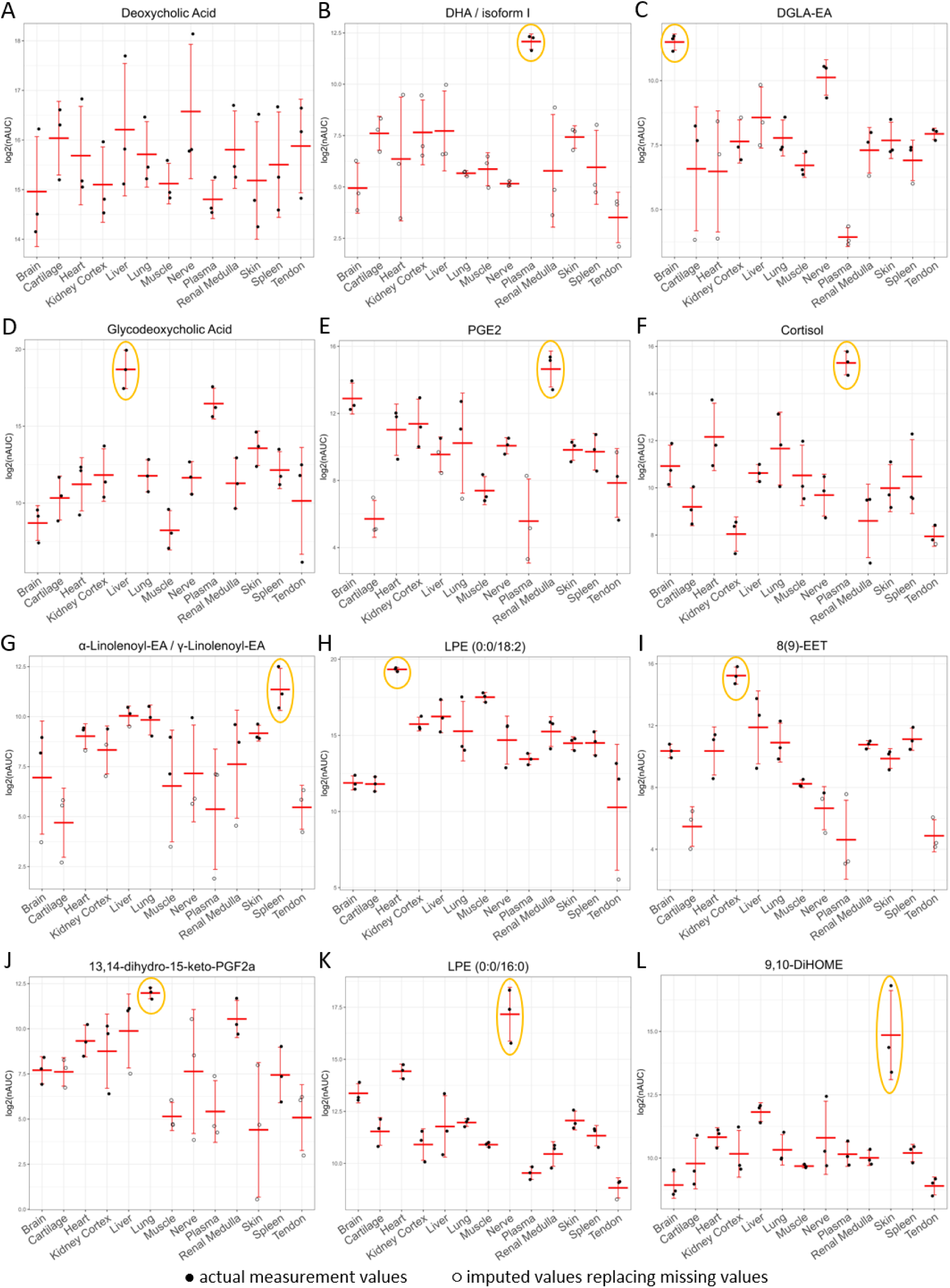
Relative abundance values, i.e. normalized area under curve (nAUC), of selected molecules showing low specificity (A: deoxycholic acid), high specificity (B: DHA isoform I, one double-bond shows trans-configuration, note that all values but for plasma are imputed) or a predominant occurrence in one organ or tissue type of C-L: DGLA-EA, Glycodeoxycholic Acid, PGE2, Cortisol, Alpha-Linolenoyl-EA/gamma-Linolenoyl-EA, LPE (0:0/18:2), 8(9)-EET, 13,14-dihydro-15-keto-PGF2alpha, LPE (0:0/16:0), and 9,10-DiHOME. Full circles, representation of the actual measurement result; open circles, imputed values replacing missing values (indicating lack of detection).

Numerous expression patterns characteristic for various tissue types were observed, as depicted in Figure 2B-L. Several of those observations are in accordance with known lipid functions as mentioned in the following. The endocannabinoid dihomo-gamma-linolenoyl ethanolamide (DGLA-EA) was predominantly found in the brain (Figure 2C) [41], the bile acid and nutrient signaling hormone glycodeoxycholic acid [42] predominantly in liver (Figure 2D), the prostaglandin (PG) and renal water transport regulator PGE2 [43] in renal medulla (Figure 2E) and the stress hormone cortisol [44] in plasma (Figure 2F). Alpha-linolenoyl ethanolamide, predominant in spleen tissue (Figure 2G), may relate to its physiological role as immune modulator for macrophages [45]. Remarkably, this analysis also revealed characteristic patterns of molecules with yet unknown functional implications. This included the lyso-phosphatidylethanolamine (LPE) (0:0/18:2) found to be predominant in the heart (Figure 2H), 8(9)-EET in the kidney cortex (Figure 2I), 13,14-dihydro-15-keto-PGF2alpha in the lung (Figure 2J), LPE (0:0/16:0) in nerve tissue (Figure 2K), and 9,10-dihydroxy-octadecenoic acid (DiHOME) in the skin (Figure 2L).

### Levels of free PUFAs and PUFA-derivatives correlate between organ types

As several immune modulatory molecules are derived from PUFAs, relative abundance values of the main PUFAs arachidonic acid (AA) and DHA were closely inspected. As demonstrated in Figure 3, high levels of PUFAs were detected in the kidney cortex, liver and brain, intermediate levels in the lung, nerve tissue, skin, spleen and renal medulla, and the lowest levels in cartilage, heart, muscle, and tendon tissues (Figure 3A and 3D). The levels of the corresponding PUFA-containing lyso-phosphatidylcholine (LPC) molecules (Figure 3B and 3E) and endocannabinoids (Figure 3C and 3F) were found to correlate with those basal PUFA levels. However, the comparative analysis demonstrated that some molecules escaped this pattern. For example, the levels of LPC (0:0/20:4) in heart tissue were higher than expected from the levels of the 20:4 fatty acid AA in heart tissue (Figure 3B), whereas LPC (0:0/22:6) in heart tissue (Figure 3E) was as close to the 22:6 fatty acid DHA (Figure 3D) as in other tissues. This specific pattern strongly suggested tissue-specific metabolic processes and motivated us to perform more systematic association analyses.

**Figure 3:**
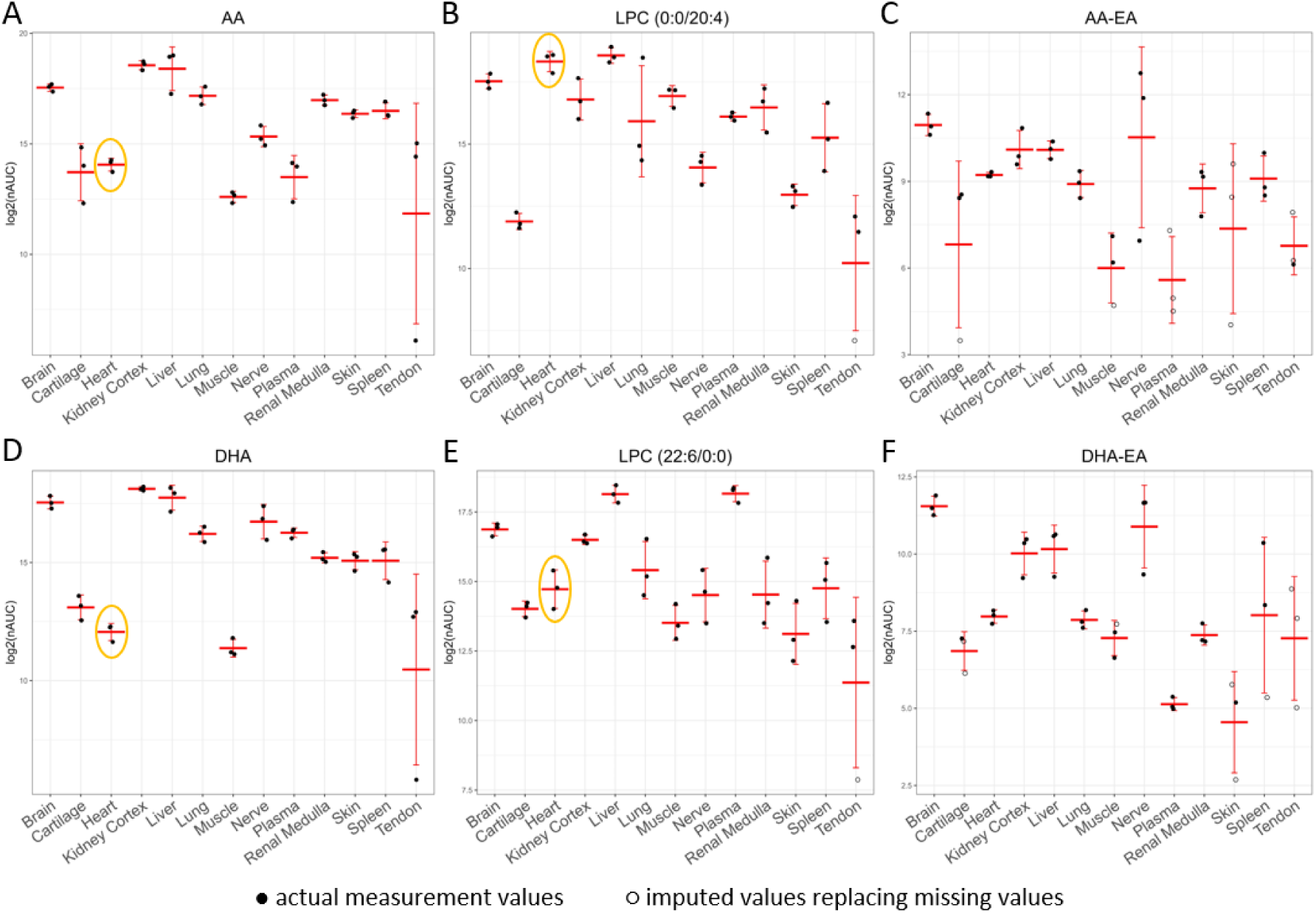
Relative abundance values, i.e., normalized area under curve (nAUC), of PUFA-related molecules derived from AA (A-C) and DHA (D-F). Note that the expression patterns of these related molecules correlate in most tissues, characteristic deviations as indicated in case of LPC (0:0/20:4) may thus indicate specific metabolic processes.

### Association analysis of lipid mediators indicates functional correlations

A heat map resulting from an association analysis of the lipid mediators detected in all tissue and organ types is displayed in Figure 4. As expected from the expression pattern depicted in Figure 3, members of the same lipid class, including fatty acids, bile acids, lysolipids, endocannabinoids and oxylipins, were found to cluster without exception (Figure 4, cluster and color-coded bar on the left-hand side). Remarkably, the oxylipin 4-HETE was found to cluster with oxylipins formed in a non-enzymatic fashion, suggesting that it may not represent an enzymatic reaction product. The anesthetic embutramide used to euthanize the sheep was included in the analysis, while it is no endogenous metabolite. However, the relative abundance differences illustrate the different properties of tissue types regarding the accumulation of this drug. Remarkably, embutramide was hardly detectable in plasma, indicating a highly efficient accumulation in the heart, lungs and brain.

**Figure 4:**
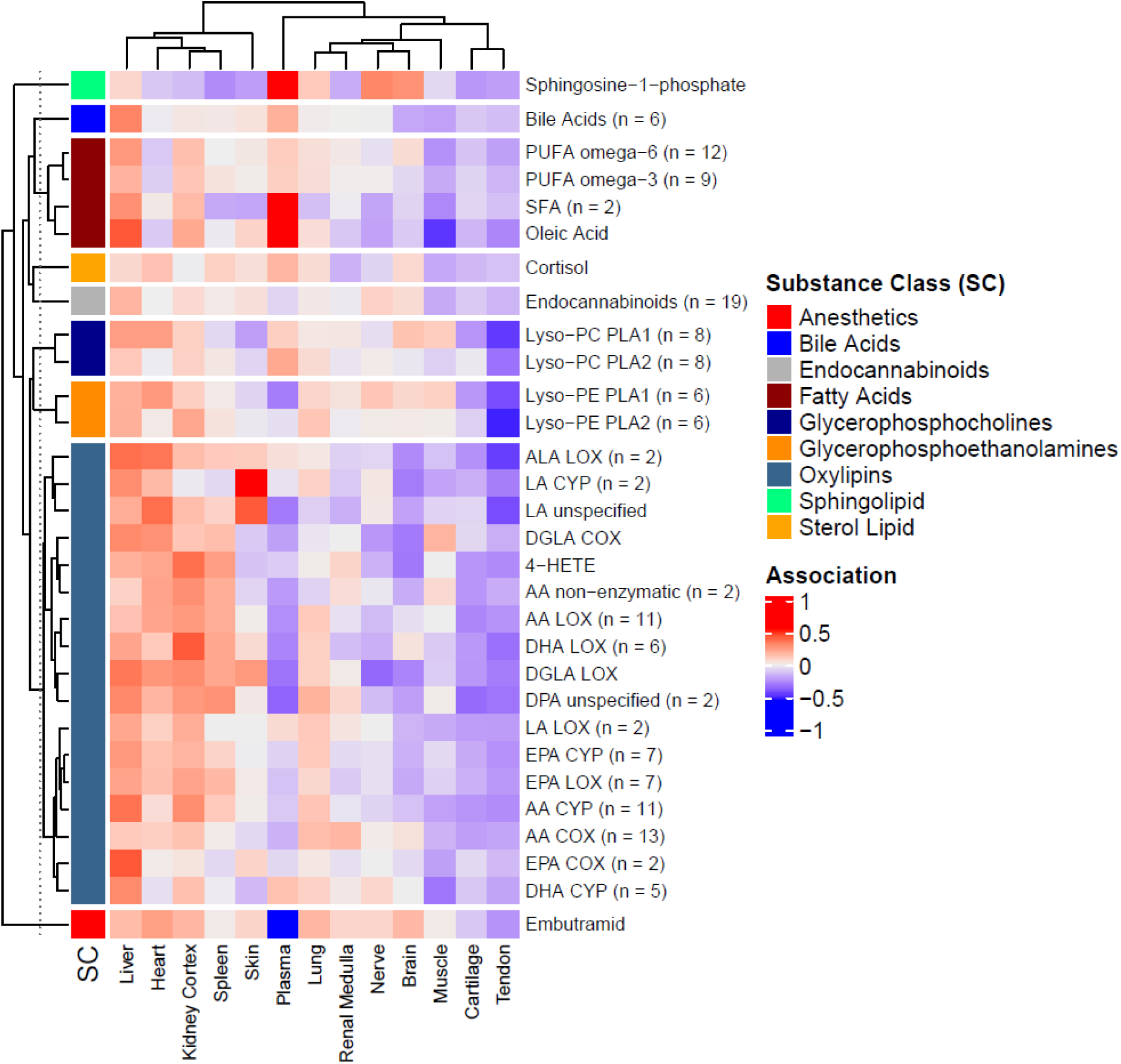
Association analysis of lipids. Hierarchical clustering grouped all oxylipins into a large cluster, oxylipin subclasses were annotated according to the precursor fatty acids: alpha-linolenic acid (ALA), linoleic acid (LA), dihomo-gamma-linolenic acid (DGLA), arachidonic acid (AA), docosahexaenoic acid (DHA), docosapentaenoic acid (DPA) and eicosapentaenoic acid (EPA); and the associated enzymes: lipoxygenase products (LOX), cytochrome p450 products (CYP), cyclooxygenase products (COX), or non-enzymatic. The previously uncharacterized oxylipin 4-HETE was found closest to oxylipins formed in a non-enzymatic fashion. The lysolipids were classified according to phospholipase A1 (PLA1) or phospholipase A2 (PLA2) products. The fatty acids were classified as saturated fatty acids (SFA), omega-3 and omega-6 polyunsaturated fatty acids (PUFAs).

The sample sources formed two main tissue cluster separated from plasma. Apparently one group represented tissue with rather high levels of lipids, in contrast to the other with rather low levels (Figure 4, cluster on the top). While heart tissue belonged to the first cluster, skeletal muscle tissue was in the second, indicating profound differences of lipid metabolism between these two muscle tissue types.

### Functional characterization of tissue types by proteome profiling

Samples from the solid tissue types and plasma were subjected to proteome profiling as described previously [28]. After filtering for reproducible and highly confident results, between 171 (cartilage) and 2761 (brain) proteins were identified per tissue type, resulting in a total of 4717 proteins covering a dynamic range of label free quantification values of 5 orders of magnitude (Supplementary Table S4). Mutual comparisons of each tissue type against the sum of all other tissue types identified proteins specifically enriched in one tissue/organ type. The numbers ranged from 461 proteins found to be enriched in brain tissue brain down to only two proteins found to be enriched in tendons (red marked proteins, Supplementary Table S4). These proteins comprised many well-known tissue-specific human orthologues as characterized by gene ontology, again qualifying this approach.

GO term enrichment analysis was used for functional characterization of organ types as depicted in Figure 5 in case of brain tissue. Processes such as “neuron projection” and “intracellular signal transduction” were found to be over-represented in brain tissue. The presynaptic membrane, synaptic vesicle and synapse were identified as hub-terms linking a large number of characteristic proteins (Figure 5). Thus, characteristic assignment of tissue-specific functions *via* the proteomics data was obtained and allowed us to perform co-correlation analyses in order to investigate potential correlations of characteristic tissue functions with lipid mediator profiles.

**Figure 5:**
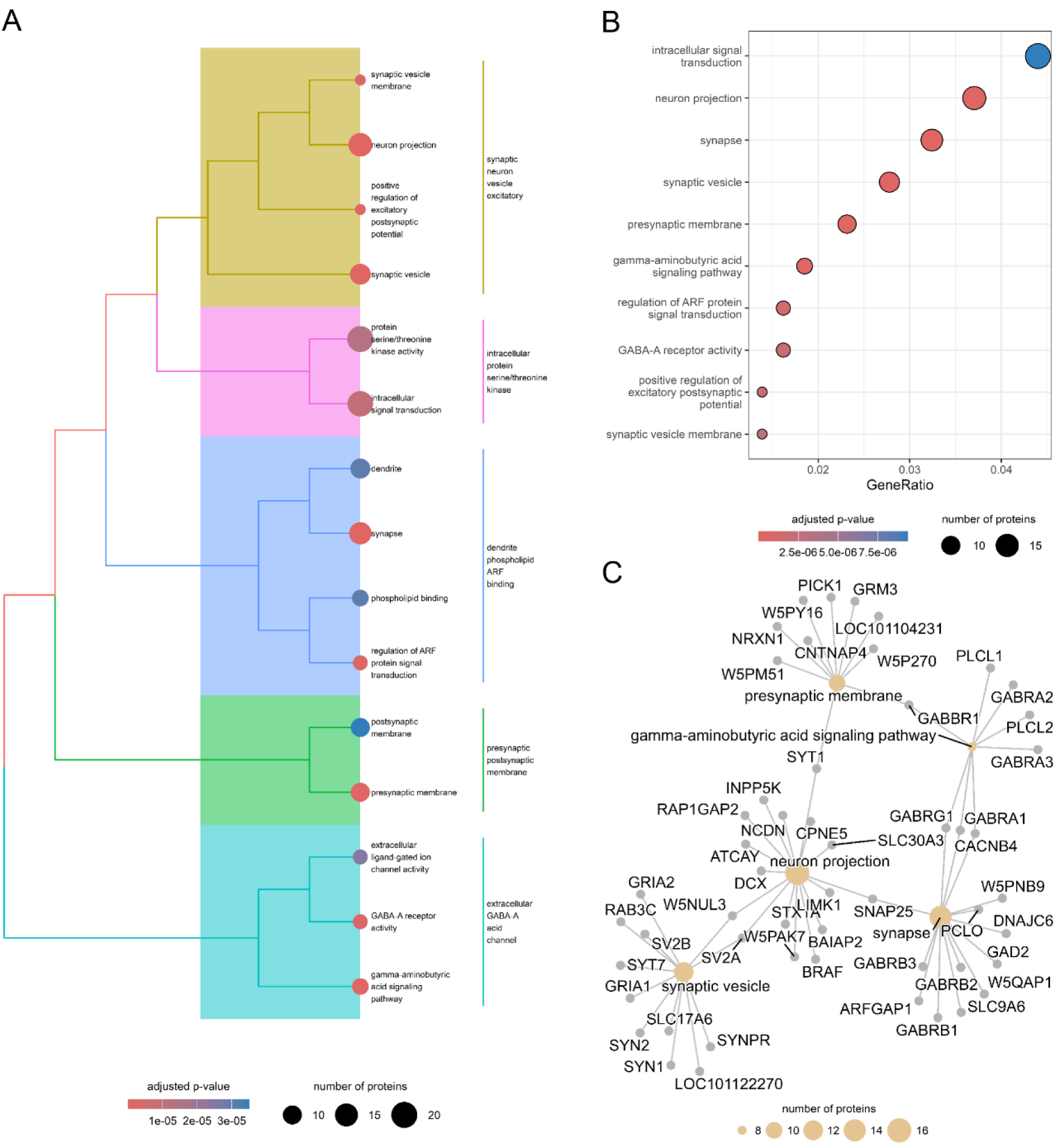
Results of the GO term enrichment analysis obtained from proteome profiling of ovine brain tissue as A) tree plot of the hierarchical cluster analysis of the enriched GO terms, B) dot plot of the 10 most significantly enriched GO terms and C) C-net plot of a network of the 5 most significantly enriched GO terms in the ovine brain tissue. Dot size indicates the number of participating proteins in the respective GO biological process. Dot color indicates the enrichment test adjusted p-value (Figure 5A and 5B).

### Co-expression analysis of small molecules and proteins reveals functional correlations

Co-expression analysis of all identified lipids with all identified proteins and clustering thereof resulted in the formation of 27 modules (Supplementary Table S5). The efficient separation of modules is demonstrated in Supplementary Figure 2 (*Supporting Information SI1*). These modules showed pronounced tissue-type specific characteristics as depicted in Figure 6. To give examples, module M12 was most representative for renal medulla, module M15 for kidney cortex, module M24 for nerve tissue. On the other hand, some modules showed characteristic under-representation, such as module M14 in case of cartilage and tendon tissue (Figure 6). While 9 modules listed proteins only, 18 modules successfully correlated proteins with small molecules, allowing for positive or negative correlations. Investigation of proteins listed in the modules by gene ontology term enrichment analysis allowed us to relate a given set of lipids with cellular functions characteristic for the given tissue type (Supplementary Table S5).

**Figure 6:**
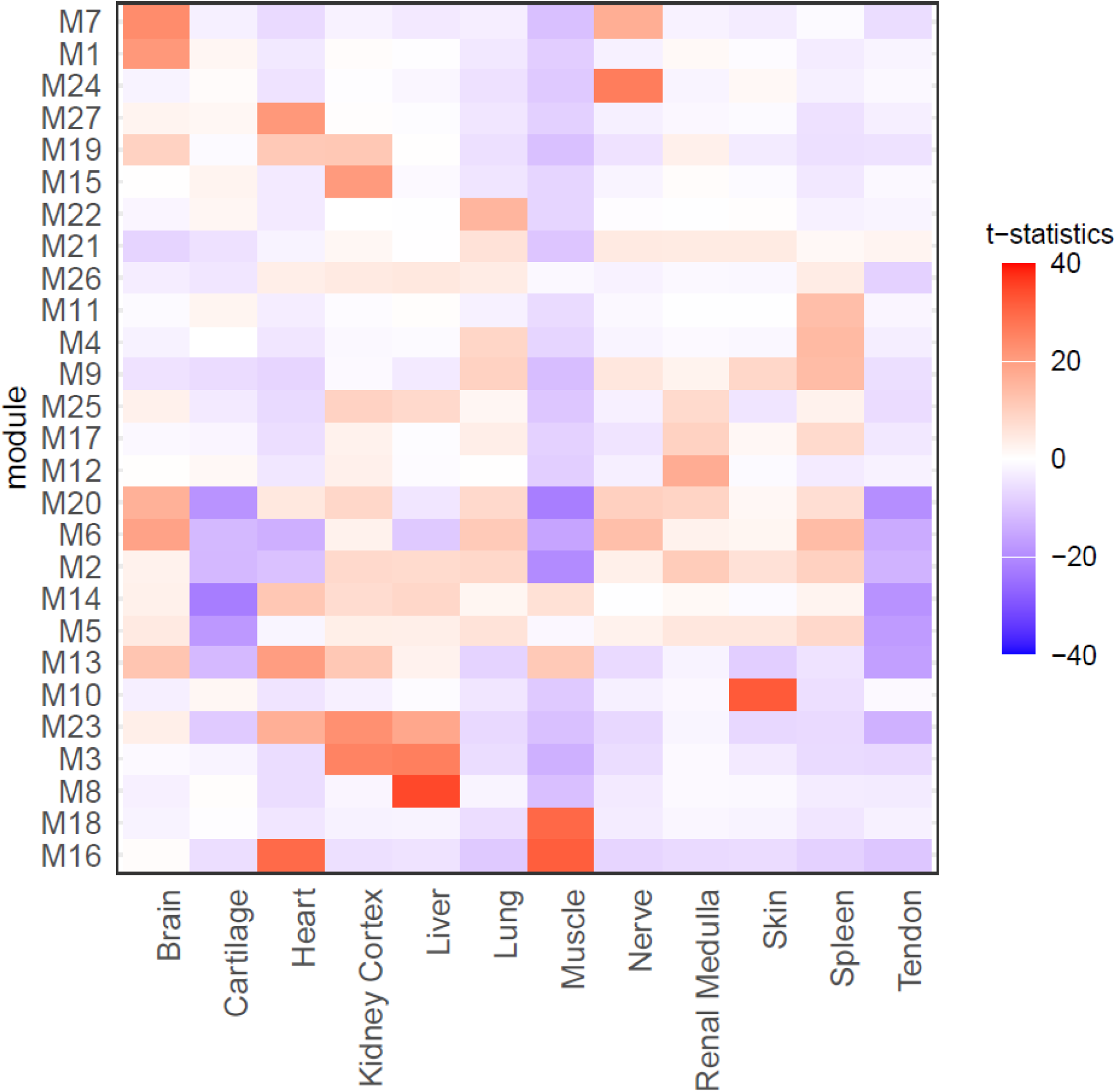
Heatmap of Eigengene analysis of the 27 modules. Several modules such as M1, M8 and M18 were found apparently specific for one organ type, others such as M3 and M16 were characteristic for two tissue types, whereas other modules such as M5 and M14 displayed characteristic negative correlations.

In order to evaluate whether the obtained results contained reliable information, the modules were screened for correlations compatible with established knowledge. Module M8 provided a positive example, correlating the CYP monooxygenase products 11,12-dihydroxy-eicosatrienoic acid (DiHETrE), 14,15-dihydroxy-eicosatetraenoic acid (DiHETE), 16,17-dihydroxy-docosapentaenoic acid (DiHDPA) and 19,20-DiHDPA, with a cluster of monooxygenases (CYP1A2, CYP2A13, CYP4F2, CYP4F3, CYP8B1, FMO1, FMO5; Supplementary Table S5). Module M8 was also found to be highly specific for the liver (Figure 6), in line with the liver-specific expression pattern of the monooxygenases [46]. Thus, module M8 successfully reproduced established biochemical knowledge.

Subsequently, the modules were evaluated with regard to potential new findings. Module M2 correlated low levels of PUFAs such as DHA, DPA, EPA, DGLA, AA and their derivatives such as 20-HDoHE with low levels of proteins involved in cytoplasmic translation and endoplasmic reticulum function. This module was characteristic for liver, lung and kidney, but showed a characteristic negative correlation in muscle, tendon and cartilage. Module M3 correlated the less unsaturated or fully saturated fatty acids oleic acid, linoleic acid and palmitic acid together with 18-HETE and 19-HETE with high levels of proteins involved in mitochondrial fatty acid beta-oxidation. Module M3 was mainly characteristic for kidney cortex and liver. Module M1, apparently specific for brain, correlated 11,12-DiHETE, 15-deoxy-PGJ2 and 5(6)-EET with proteins mediating chemical synaptic transmission. Module M18 correlated arachidonoyl ethanolamide phosphate with proteins essential for muscle contraction. Module M26 correlated numerous LPCs and LPEs together with several oxylipins with proteins characteristic for extracellular exosomes.

Other modules clustering proteins with characteristic functions, such as module M17 associated with mRNA splicing, module M21 associated with extracellular matrix components and module M22 associated with angiogenesis, did not show any significant correlations with lipids.

## Discussion

Here we present a tissue atlas of lipid mediators comprising fatty acids, oxylipins, endocannabinoids and bile acids in combination with a corresponding proteome atlas of twelve different tissues and organs and plasma from healthy adult sheep, *Ovis aries*. While no tissue-specific lipid was identified such as the numerous tissue-specific proteins, lipid patterns characteristic for tissues were clearly discerned (Figure 2). To the best of our knowledge, this is the first study systematically investigating tissue and organ expression characteristics of lipid mediators in relation to proteins. By carefully sampling freshly isolated tissue samples from euthanized animals, we were able to obtain reliable expression pattern of molecules which may be associated with methodological obstacles [47]. We have previously described the deregulation of eicosanoids and other lipid mediators in the context of chronic inflammation [48], ovarian cancer [49] and heart diseases such as functional mitral regurgitation [50] and clearly demonstrated the power of oxylipins at nanomolar concentrations for the modulation of inflammatory processes in cell culture experiments [51]. Using a data-dependent acquisition method, we were able to identify the previously undefined oxylipin 4-HETE (Figure 1). The reliable identification of this arachidonic acid (AA)-derived compound was verified *via* chemical synthesis. While this compound was associated with non-enzymatically formed AA products in the hierarchical clustering (Figure 4), it is most plausible that it may be formed *via* an allylic hydroxylation by CYP *in vivo* since AA lacks a double bond between C4 and C5 for potential autoxidation [52].

Clustering the tissue types according to lipid profiles separated three main tissue types according to levels of lipid mediators (Figure 4). Unexpectedly, heart tissue was clustered with tissues showing higher levels of lipid mediators, while skeletal muscle was in the other tissue cluster with low levels of lipid mediators, indicating profound differences in the lipid metabolism of these muscle tissues (Figure 4). Indeed, heart tissue was described to have high levels of lipoprotein lipase in contrast to skeletal muscle [53], which apparently accounts for specific metabolic properties of these tissues and provides a biochemical explanation for the present observation.

The anesthetic embutramide used to euthanize the sheep was included in the analysis, while it is no endogenous metabolite. However, the relative abundance differences illustrate the dynamics of redistribution of a pharmacologic substance relating to the different properties of tissue types. Figure 4 illustrates tissue-specific uptake of this drug, a highly efficient uptake in the heart, lungs and brain in contrast to poor uptake in muscle, tendon and cartilage.

Most lipids were detected in almost all samples, indicating their ubiquitous presence. However, the lipid composition of plasma was found to differ profoundly from solid organs and tissues (Figure 4). This may not be surprising, as plasma lipids mainly derive from lipoprotein particles, whereas tissue lipids mainly derive from cell membranes and lipid stores. As many lipid mediators have been described to be rather short-lived [54], plasma levels of such lipids quickly fall below detection thresholds. This seems to apply e.g. to the prostaglandin PGE2, described to be almost completely degraded within one minute in plasma [55]. In this study, PGE2 was identified in all tissue types but cartilage and plasma (Figure 2), indicating that the presently applied sample preparation protocol allowed us to preserve meaningful molecules while being short-lived. The short half-life of PGE2 and the substantial amounts detected in the kidney cortex, brain, nerve and heart tissue indicate that continuous synthesis takes place in the healthy organs which may indicate physiological relevance such as described for water transport in renal collecting ducts [43]. In a previous study, PGE2 was detected in plasma derived from human patients suffering from acute inflammation, but not in healthy controls [27]. This is in line with the present observation. We thus suggest that several other lipid mediators presently detected in organs and tissues may have relevant physiological functions.

In order to learn more about such physiological functions, we have conducted a correlation analysis of lipids with proteins making use of the ample functional annotation of proteins *via* gene ontology [17]. As most lipids were detected at varying amounts in the different organ and tissue types, we expected meaningful functional correlations. Indeed, this approach delivered correlations which are obviously meaningful, such as the preferential formation of monooxygenase products in liver tissue rich in the corresponding enzymes, the monooxygenases. In addition, the observed pattern of 18-HETE and 19-HETE with proteins involved in mitochondrial beta-oxidation, characteristic for the kidney cortex (module M3 of Figure 6) is fully consistent with established knowledge. 19-HETE has been described to be essential for the stimulation of proximal renal tubules responsible for the reabsorption of water, ions and nutrients – an energy-consuming process enabled through the oxidative metabolism of the high number of mitochondria present in these tubules [56; 57]. The Eigengene analysis depicted in Figure 6 indicated a correlation between the oxylipins 11,12-DiHETE, 15-deoxy-PGJ2 and 5(6)-EET and chemical synaptic transmission in the ovine brain (module M1, Figure 6), another striking correlation. These oxylipins have indeed been described to occur in the brain or the central nervous system [58–60]. While 5(6)-EET seems to be directly involved in synaptic transmission [61], 15-deoxy-PGJ2 is a potent modulator of this physiological process [62].

A potentially new finding is the presently observed correlation of PUFAs with endoplasmic reticulum and ribosomal proteins, all of which are involved in protein synthesis. A lot of literature exists with regard to functional implications of PUFAs. High levels of PUFAs seem to support the function of the blood-brain barrier and cardiovascular health, potentially due to anti-inflammatory properties [63; 64]. The high content of PUFAs in cardiolipins required for appropriate mitochondrial functions was described to point to a correlation of PUFAs with mitochondrial content [65]. However, the present data demonstrate that high levels of mitochondrial proteins in muscle and heart were associated with low levels of PUFAs, not supporting this assumption. Apparently, PUFAs were found to be most abundant in three tissue types, namely in brain, liver and kidney cortex (Figure 3). This observation hardly supports the previously suggested functional associations. Neither longevity as attributed to brain cells, nor the presence of immune-modulatory cells as evident in case of the spleen or the high mitochondrial content as evident for muscle tissue is characteristic for these three tissue types. However, high levels of membrane-rich endoplasmic reticulum are indeed characteristic for cells with high protein synthesis and account for the morphological appearance of e.g., plasma cells. The chaperoning function of the endoplasmic reticulum (ER) has been described to be critically dependent on lipids showing a broad range of chemical diversity, thus also rich in PUFAs [66]. A specific support of chaperoning functions by PUFA-containing endoplasmic reticulum membrane lipids may thus explain the present observation, which is fully compatible with previous observations and thus represents a new model for the functional role of PUFAs with intriguing implications.

Tissue types with little protein synthesis and turnover such as cartilage, muscle and tendon were found to contain low levels of PUFAs and the corresponding, potentially anti-inflammatory oxylipins, designated as pro-resolving mediators [67]. Indeed, pro-resolving mediators have been described to mediate efferocytosis, the removal of dead cells and debris [68]. A lack of omega-3 fatty acid-derived lipid mediators, apparently also associated with ageing, may thus limit efferocytosis and drive chronic inflammation [69; 70]. As a consequence, it makes sense to relate low levels of pro-resolving mediators in cartilage, muscle and tendon with an inherent risk for chronic inflammatory diseases such as osteoarthritis or tendinopathies. Actually chronic inflammation is a characteristic feature of most age-associated health threats including cardiovascular diseases, diabetes, osteoarthritis, cancer and neurodegenerative diseases [71]. The increased risk for such diseases with age seems also to be associated with decreased protein and lipid turnover [72; 73], and increased protein misfolding [74], eventually resulting in inflammasome activation and sustained inflammation [75].

Protein chaperoning and replacement of damaged protein *via* synthesis may thus synergize with anti-inflammatory properties of PUFA derivatives, eventually promoting the resolution of inflammation. Evidently, an improved understanding of the biochemistry of lipid mediators at physiologic conditions will support the design of therapeutic strategies to promote resolution pathways [76–78].

### Limitations of the study

This is a feasibility study. Thus, the statistical power of this study is limited. The results represent correlations that were interpreted according to established biochemical mechanisms, but definite causal evidence is still lacking. The overall resolution of the analyses will need further improvement in order to gain more accurate annotations.

### Conclusions

To the best of our knowledge, this is the first co-correlation study of oxylipins with proteins across different tissue types. The aim of the study was to project functional knowledge regarding proteins to correlate fatty acids and oxylipins, for which such knowledge is largely missing. While this remains to be a feasibility study, the results allowed us to validate the approach *via* the independent confirmation of previous knowledge. Intriguingly, the results also suggest a new model that could explain the strong empirical correlation between a lack of omega-3 fatty acids and other PUFAs and higher risk for chronic inflammation. The data suggested that high levels of PUFAs may be essential for proper function of the cellular protein synthesis and folding machinery, in addition to their role as precursors of pro-resolving mediators. This new model for the physiologic function of PUFAs offers far-reaching biomedical implications. It represents a testable hypothesis, that can be rigorously challenged by experiments designed specifically for this purpose.

## Supporting information

**Supporting Information SI1**: Detailed description of proteomics and lipidomics sample preparation, data acquisition and processing, and of the synthesis of 4-HETE and the GC-MS analysis of the standard. Supplementary Figure 1 – MS2 spectrum of *m/z* = 343.2279 representing 4-HDoHE. Supplementary Figure 2 – Separation efficiency of the modules.

**Supplementary Table S1:** Internal standards used for the lipidomics analysis

**Supplementary Table S2**: Degree of identification of the lipids detected in sheep organ and tissue samples.

**Supplementary Table S3**: m/z values specific for oxylipins and their precursor fatty acids used for the inclusion list

**Supplementary Table S4**: List of proteins identified in sheep organ and tissue samples. Proteins identified in only organ or tissue type, but not in any other, are marked in red. A total of 4717 proteins was identified (FDR<0.01 on peptide and protein level). Proteins reproducibly identified (in at least 4 of 6 replicates) in one tissue/organ type but no other are marked in red: brain (461), heart (26), kidney cortex (102), renal medulla (63), liver (102), lung (66), muscle (60), nerve (33), skin (88), spleen (218), tendon (2), cartilage (15).

**Supplementary Table S5**: List of all 27 modules identified. The uniprot accession numbers of the human orthologs as applied for the cluster and co-expression analysis are listed. 18 modules contain proteins and lipids, 9 modules contain only proteins. As depicted in Figure 6, the Eigengene analysis suggests positive or negative associations with tissue/organ types. Gene set enrichment analysis of the proteins suggests gene ontology (GO) terms significantly enriched (adjusted p-values according to Benjamini-Hochberg).

## Acknowledgements

The authors are grateful to the Core Facility of Mass Spectrometry at the Faculty of Chemistry and the Joint Metabolome Facility, both members of the Vienna Life-Science Instruments (VLSI).

