## Supplementary Information SI1 for "Integrated tissue proteomics and lipidomics from ovine organs suggests novel functional implications for fatty acids and lipid mediators"

**Integrated lipid mediator and proteome profiling links polyunsaturated fatty acids to protein folding functions**

^*^ contributed equally

Correspondence:

### Materials and methods

#### Proteomics Sample Preparation

For untargeted tissue and plasma proteomics analysis, frozen tissue samples were transferred into a 15 mL Falcon^TM^ tube containing 150 µL of lysis buffer (8 M urea, 50 mM triethylammonium bicarbonate (TEAB), 5 % sodium dodecyl sulfate (SDS)) and homogenized using an ultrasound sonication homogenizer. After centrifugation at 5000 g and room temperature (RT) for 5 min, the supernatant was transferred into a 1.5 mL Eppendorf microcentrifuge tube. Ovine plasma was diluted 1:20 using lysis buffer, and tissue and plasma samples were heated for 5 min at 95 °C and 300 rpm under constant shaking. After cooling to RT, the protein concentration was determined *via* a bicinchoninic acid (BCA) assay. Aliquots of plasma and tissue samples containing 20 µg protein were used for enzymatic protein digestion according to the ProtiFi S-Trap^TM^ protocol with some modifications. In short, the proteins of diluted samples (20 µg of protein in 20 µL lysis buffer) were reduced and carbamidomethylated by adding 64 mM dithiothreitol (DTT) and 486 mM iodoacetamide (IAA). After addition of S-Trap buffer (90 % v/v methanol, 0.1 M triethylammonium bicarbonate), the samples were loaded onto S-trap mini cartridges. The samples were washed 4 times with S-trap buffer and subsequently digested using Trypsin/Lys-C Mix (MS grade; Promega Corporation, Madison, WI, USA) in a ratio of 1:40 at 37 °C for two hours. The eluted peptides were dried *via* vacuum concentration and stored at -20 °C until LC-MS analysis.

#### Proteomics Data Acquisition

For reconstitution of peptides, 5 µL 30 % formic acid (FA), containing four synthetic peptides for quality control, were added to the dried samples. Samples were further diluted with 40 µL of loading solvent (97.9 % H_2_O, 2 % acetonitrile (ACN), 0.05 % trifluoroacetic acid (TFA)). Separation and analysis of peptides was achieved using a Dionex UltiMate 3000 Nano LC system (Thermo Fisher Scientific) hyphenated to a timsTOF Pro mass spectrometer (Bruker). Pre-concentration of peptides was performed *via* a pre-column (2 cm x 100 µm, 5 µm, 100 Å, C18 Acclaim PepMap^TM^ 100, Thermo Fisher Scientific) operating at a flow rate of 10 µL min^-1^ using mobile A (99.9 % H_2_O, 0.1 % FA). For separation of peptides, a captive spray compatible reversed-phase analytical column (25 cm x 75 µm, 1.6 µm, 120 Å, C18 Aurora Series emitter column, IonOpticks) was used. Here, a gradient flow profile was applied starting at 7 % mobile phase B (79.9 % ACN, 20 % H_2_O, 0.1 % FA) and going up to 40 % B over 95 min or over 43 min for tissue and plasma samples, respectively, at a flow rate of 300 nL min^-1^, resulting in a total chromatographic run time of 135 min or 85 min, respectively, containing washing and equilibration steps. For the mass spectrometric analysis with the timsTOF Pro, the captive spray ion source was operating at 1650 V in the Parallel Accumulation-Serial Fragmentation (PASEF) mode applying moderate MS data reduction. The scan range was 100 to 1700 m/z for MS and MS/MS spectra recording and the 1/k0 scan range was set to 0.60 – 1.60 V*s cm^-2^ resulting in a ramp time of 100 ms to achieve trapped ion mobility separation.

#### Proteomics Data Processing

Protein identification was performed *via* MaxQuant (version 1.6.17.0) employing the Andromeda search engine against the UniProt Database (version 11/2021, 20’375 entries). A mass tolerance of 20 ppm for MS spectra and 40 ppm for MS/MS spectra, a PSM–, protein– and site-false discovery rate (FDR) of 0.01 and a maximum of two missed cleavages per peptide were allowed. Match-between-runs was enabled with a matching time window of 0.7 min and an alignment time window of 20 min. Oxidation of methionine, N-terminal protein acetylation and phosphorylation of serine, threonine and tyrosine were set as variable modifications. Carbamidomethylation of cysteine was set as fixed modification. Global proteome data analysis was performed *via* Perseus (version 1.6.14.0). Proteins with at least 60 % identification rate in at least one group were considered for analysis. A two-sided Student’s t-test with S0 = 0.1 and an FDR cut-off of 0.05 was applied for identifying multiple testing-corrected significantly regulated proteins.

#### Lipidomics Sample Preparation

For untargeted plasma analysis of oxylipins and fatty acids, EDTA-anticoagulated plasma was thawed on ice and 400 mg of plasma was added to a 15 mL Falcon^TM^ tube containing ice-cold EtOH (1.6 mL, abs. 99.9 %, AustrAlco) and an internal standard mixture (Supplementary Table S1). All standards, including those indicated in Supplementary Table S2, were purchased by Cayman Chemical, Michigan, USA. The suspension was vortex mixed and stored at -20 °C overnight for protein precipitation. For tissue oxylipins analysis, the Falcon^TM^ tubes containing the tissue samples and 1 mL of EtOH were transferred from -20 °C to ice and samples were homogenized using an ultrasound sonication homogenizer and stored at -20 °C overnight. Then, 500 µL of LC-MS grade H_2_O and an internal standard mixture with the same internal standards and concentrations as used for the plasma analysis was added to each sample. Subsequently, plasma and tissue samples were centrifuged (30 min, 4536 g, 4 °C) and the supernatant was transferred into a new 15 mL Falcon^TM^ tube. To restore the original sample volume, EtOH was evaporated *via* vacuum centrifugation at 37 °C. StrataX solid phase extraction (SPE) columns (30 mg mL^-1^, Phenomenex, Torrance, CA, USA) were preconditioned with LC-MS grade methanol (2 mL; MeOH; VWR International, Vienna, Austria) and LC-MS grade H_2_O (2 mL; VWR International, Vienna, Austria). Samples were then loaded using Pasteur pipettes, SPE columns were washed with ice-cold LC-MS grade H_2_O (5 mL; VWR International, Vienna, Austria) and analytes were eluted with ice-cold LC-MS grade MeOH (500 µL; VWR International, Vienna, Austria) including 2 % formic acid (FA; ≥ 99 %; VWR International, Vienna, Austria). The eluted samples were dried using a gentle nitrogen stream at room temperature and the dried samples were reconstituted with 150 µL reconstitution solvent (H_2_O/ACN/MeOH + 0.2 % FA – vol% 65:31.5:3.5). The reconstituted samples were subsequently transferred into an autosampler held at 4 °C and measured *via* LC-MS/MS.

#### Lipidomics Data Acquisition

For the LC-MS/MS analyses, a Vanquish™ UHPLC system (Thermo Fisher Scientific™, Vienna, Austria) was coupled to a high-resolution quadrupole orbitrap mass spectrometer (Thermo Fisher Scientific™ Q Exactive™ HF hybrid quadrupole orbitrap mass spectrometer). Therefore, the Vanquish™ ultra-high-performance LC (UHPLC) system was equipped with a reversed-phase Kinetex^®^ XB-C18 column (2.6 µm XB-C18, 100 Å, LC column 150 × 2.1 mm, Torrance, CA, USA) to achieve analyte separation. The flow rate was set to 200 µL min^-1^ and the injection volume was set to 20 µL. All samples were measured in technical duplicates in negative mode and measured once in positive ion mode. Blank injections were performed after each sample and its technical duplicate to monitor potential carryover and ensure the reliability of the analytical results. The LC column oven was set to 40 °C and the autosampler temperature was held at 4 °C. A gradient flow profile was applied starting at 35 % B with mobile phase A representing H_2_O + 0.2 % FA and mobile phase B representing ACN:MeOH (vol% 90:10) + 0.2 % FA. Solvent B was increased to 90 % B (1-10 min), further increased to 99 % B within 0.5 min and hold for 5 min. After decreasing to 35 % B within 0.5 min, the column was equilibrated for 4 min, resulting in a total run time of 20 min. The Q Exactive™ HF high-resolution mass spectrometer was equipped with a HESI source operating in negative and positive mode with spray voltage set to 3.5 kV in both modes, capillary temperature set to 253 °C in negative mode and 300 °C in positive mode, sheath gas and auxiliary gas set to 46 and 10 arbitrary units in both modes, respectively. Data were recorded in the scan range of 250-700 m/z on the MS1 level with a resolution of 60,000 (*m/z* 200). For the MS/MS fragmentation in the data-dependent acquisition (DDA) mode, a Top 2 method with a resolution of 15,000 (*m/z* 200) was applied using an HCD collision cell with a normalized collision energy of 24. Here, 33 *m/z* values specific for oxylipins and their precursor fatty acids from an inclusion list were preferentially selected for fragmentation (Supplementary Table S3).

#### Lipidomics Data Processing

For data analysis, raw files generated by the Q Exactive™ HF high-resolution mass spectrometer were evaluated using the TraceFinder software (version 4.1; Thermo Fisher Scientific™, Vienna, Austria) making use of measurements of purchased chemical standards (Supplementary Table S2). Additionally, MS/MS fragmentation spectra were evaluated referring to reference spectra of the commercially available standards or to reference spectra from the LIPID MAPS repository library [35] using the Xcalibur™ Qual Browser software (version 4.1.31.9; Thermo Fisher Scientific™, Vienna, Austria). Coeluting isomers were reported giving the names of both analytes, e.g. 12-HETE/8-HETE, as suggested by the oxylipin community [36]. The TraceFinder software (version 4.1; Thermo Fisher Scientific™, Vienna, Austria) was used for relative quantification allowing a mass deviation of 5 ppm. The resulting peak areas of each analyte with a S/N greater than 10 were exported and read using the R software package (version 4.2.0) [37]. The peak areas were log2-transformed and analyte peak areas were normalized to the internal standards to correct for variances arising from sample extraction and LC-MS/MS analysis. Therefore, the log2-transformed mean peak area of the internal standards was subtracted from the log2-transformed analyte areas. The peak areas of the ovine plasma, organ and tissue samples were also normalized to the weight. Only molecules independently identified in all replicates of at least one tissue type were included in the statistical analysis. As several molecules turned out to occur at the limit of detection in other tissue types, data imputation was used to enable subsequent statistical analysis. To enable an imputation of missing values using the minProb function of the imputeLCMD package (version 2.1), 20 was added to the log2-transformed normalized areas to obtain values similar to proteomics label free quantification (LFQ) values.

#### Synthesis of 4(*R,S*)-hydroxy-5(*Z*),8(*Z*),11(*Z*),14(*Z*)-eicosatetraenoic acid (4-HETE)

The title compound was prepared by coupling of methyl 4-hydroxy-5-hexynoate to 1-bromo-2,5,8-tetradecatriyne followed by semihydrogenation and alkaline hydrolysis.

*Methyl 4-hydroxy-5-hexynoate (****4****)* - Ethynyltrimethylsilane (**1**, 11.71 mmol) and methyl 4-chloro-4-oxobutyrate (**2**; 11.15 mmol), both purchased from Merck Co., in 12.5 mL of dry dichloromethane were added dropwise under stirring at 0^o^C to dry AlCl_3_ (33.5 mmol) suspended in 50 mL of dichloromethane. The mixture was warmed to 22^o^C and stirred at this temperature for 3 h (*cf.* ref. 1). After cooling to 0^o^C, 1 M HCl was added and the solution extracted with diethyl ether. The product was subjected to silica gel open column chromatography affording essentially pure **3** as a light-yellow oil (9.7 mmol, yield 87%) (Scheme). This material was dissolved in 41 mL of methanol, cooled to -20^o^C, and NaBH_4_ (3.4 mmol) was added under stirring. The temperature was allowed to rise to 0^o^C under a period of 2 h after which time the mixture was stirred at 22^o^C for 64 h. Saturated NH_4_Cl solution was added, and the mixture was extracted with two portions of diethyl ether. Following purification on a silica gel column the title compound was obtained as a colorless oil (6.8 mmol, overall yield, 61%). GC-MS analysis of this material (Me_3_Si ether derivative) gave a single peak and a mass spectrum showing prominent ions at *e.g. m/z* 199 (M^+^ - CH_3_), 183 (M^+^ - OCH_3_), 167 (M^+^ - (CH_3_ + OCH_3_)), 127 (M^+^ - CH_2_-CH_2_-COOCH_3_), 89 (Me_3_SiO^+^), and 73 (Me_3_Si^+^).

*1-Bromo-2,5,8-tetradecatriyne (****7****)* - The tetrahydropyranyloxy derivative of 2,5-hexadiyn-1-ol (**5**, 10 mmol) and 1-bromo-2-octyne (**6**, 10 mmol) in 28 mL of dry dimethyl formamide were stirred at 22^o^C for 2 h in the presence of CuI (10 mmol), Cs_2_CO_3_ (20 mmol), and NaI (20 mmol) (*cf.* ref. 2). Quenching by addition of NH_4_Cl and extractive isolation afforded a product which was directly brominated using triphenylphosphonium bromide (13 mmol) in 75 mL of dichloromethane (3). The title compound obtained after silica gel chromatography was a light-brown oil (8.3 mmol, purity >90%, yield of pure compound, >75%) showing *m/z* 264 and 266 (M^+^), 155, 141 and 128.

*Methyl 4-hydroxy-5,8,11,14-eicosatetraynoate (****8****)* - Methyl 4-hydroxy-5-hexynoate (**4**, 5.3 mmol) and 1-bromo-2,5,8-tetradecatriyne (**7**, 5.7 mmol) in 20 mL of dimethylformamide were stirred at 22^o^C for 2 h with CuI (5.3 mmol), Cs_2_CO_3_ (10.6 mmol), and NaI (10.6 mmol). Purification by silica gel chromatography afforded the title compound (2.85 mmol, purity >98%, yield, 54%) as an essentially colorless oil. The mass spectrum of the Me_3_Si derivative showed *e.g. m/z* 383 (M^+^ - CH_3_), 311 (M^+^ - CH_2_-CH_2_-COOCH_3_), 213, 189 (Me_3_SiO^+^=CH-CH_2_-CH_2_-COOCH_3_), and 165.

*4-Hydroxy-5,8,11,14-eicosatetraenoic acid (****9****)* - Methyl 4-hydroxy-5,8,11,14-eicosatetraynoate (**8**, 2.8 mmol) in 100 mL of methanol was semihydrogenated by stirring under hydrogen gas for 3 h with 250 mg of Lindlar catalyst (ChemScene, Monmouth Junction, NJ) and 0.4 mL of quinoline. The product contained significant amounts of unreacted monoyne and other products and was purified by silica gel chromatography followed by reversed-phase HPLC using a 250 x 10 mm column of Nucleosil 100-7 C_18_ and a solvent system of CH_3_CN-H_2_O (80:20, v/v) at a flow rate of 4 mL/min. Saponification by treatment with 0.4 M NaOH in 80% ethanol at 22^o^C for 15 h followed by careful acidification to pH 3 and extraction with ethyl acetate afforded the free acid (purity, >95%, 0.56 mmol, yield from tetrayne, 20%). Part of this material was subjected to final purification using normal-phase HPLC using a 250 x 10 mm column of Nucleosil 50-7. The solvent system was 2-propanol-hexane-acetic acid (2.5:97.5:0.01, v/v/v) at a flow rate of 4 mL/min. This removed a trace of the presumed -lactone form of **9** (showing *m/z* 302 (M^+^)) and afforded the title compound as a colorless oil (purity, >98%). The UV spectrum of this material showed only end-absorption thus excluding the presence of impurities having conjugated double bonds. GC-MS analysis of the Me_3_Si ether/ester derivative showed a single peak giving *e.g. m/z* 464 (M^+^), 449 (M^+^ - CH_3_), 374 (M^+^ - Me_3_SiOH), 319 (M^+^ - CH_2_-CH_2_-COOSiMe_3_), 286 (-cleavage at C-7/C-8, *i.e*. loss of C-8 to C-20 plus 1 H. This is an interesting fragmentation which has previously been observed *e.g.* in the mass spectra of the Me_3_Si derivatives of 8- and 14-hydroxylinoleates (4,5)), 273 ([CH=CH-CH(OSiMe_3_)-CH_2_-CH_2_-COOSiMe_3_]^+^ or its equivalent), and 247 (Me_3_SiO^+^=CH-CH_2_-CH_2_-COOSiMe_3_).

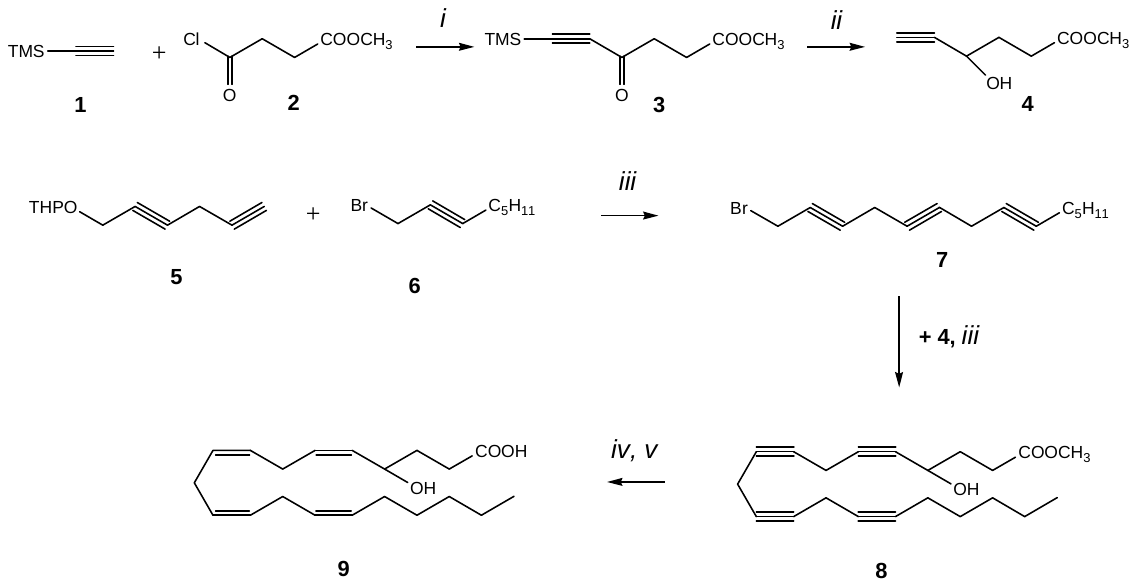

**Scheme:** Outline of synthesis of 4-HETE. *i*, AlCl_3_; *ii*, NaBH_4_; *iii*, CuI, Cs_2_CO_3_, NaI; *iv*, H_2_, Lindlar catalyst, quinoline; *v*, NaOH in aq. ethanol. "TMS", trimethylsilyl; "THPO", tetrahydropyranyloxy.

### Supplementary Figures

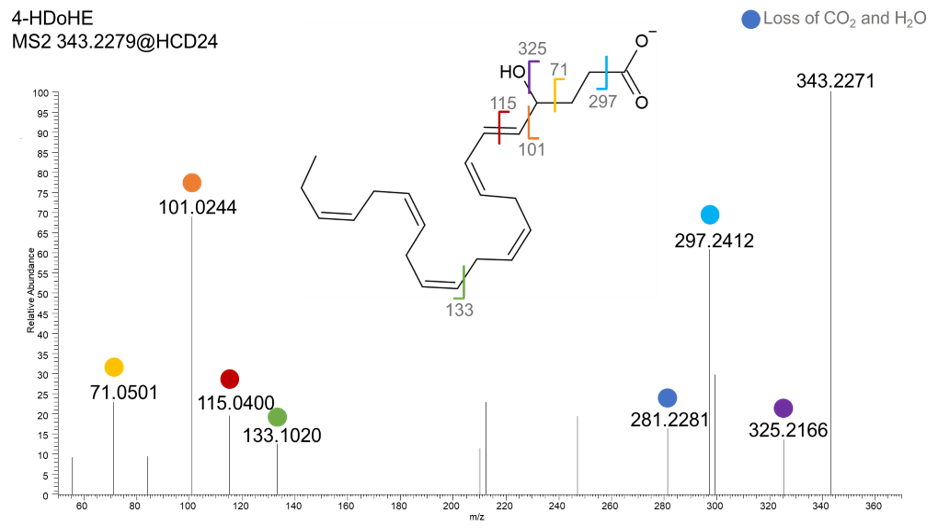

Supplementary Figure 1: MS2 spectrum of *m/z* = 343.2279 representing 4-HDoHE

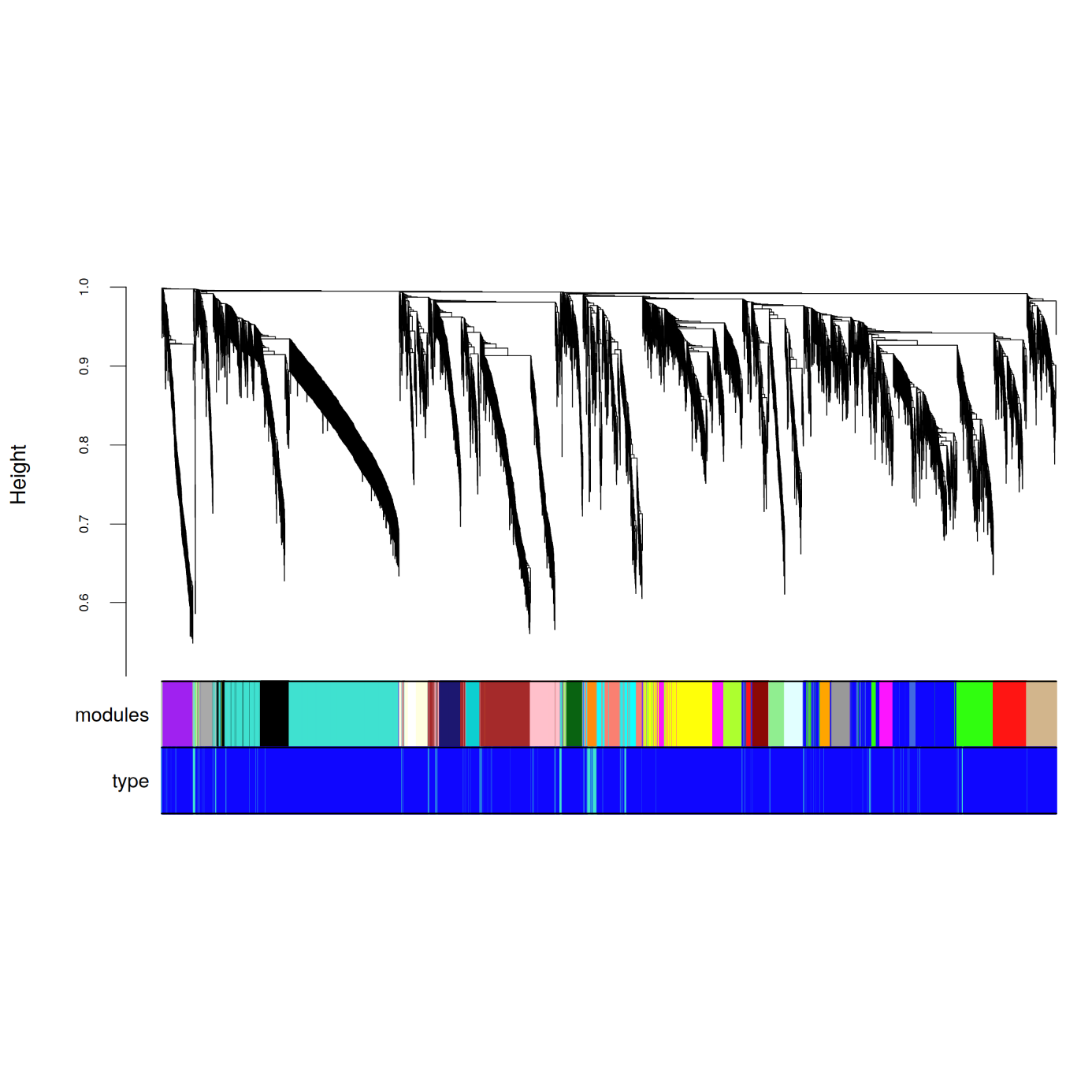

Supplementary Figure 2: Separation efficiency of the modules
